# Response to Tavares et al., “Covariation analysis with improved parameters reveals conservation in lncRNA structures”

**DOI:** 10.1101/2020.02.18.955047

**Authors:** Elena Rivas, Sean R. Eddy

## Abstract

Tavares’ conclusions depend on an assumption that the statistic they use (RAFS) is an appropriate measure of RNA base pair covariation, but RAFS was not designed to measure covariation alone. RAFS detects positive signals in common patterns of primary sequence conservation in absence of any covariation. To illustrate the severity of the problem, we show that Tavares’ analysis reports “significantly covarying base pairs” in 100% identical sequence alignments with no variation or covariation. We use Tavares’ sequence alignment of HOTAIR domain 1 as an example to show that the base pairs they identify as significantly covarying actually arise from primary sequence conservation patterns. Their analysis still reports similar numbers of “significant covarying” base pairs in a negative control in which we permute residues in independent alignment columns to destroy covariation. There remains no significant covariation support for evolutionarily conserved RNA structure in the HOTAIR lncRNA or other lncRNA structures and alignments we have analyzed.

---

A recent bioRxiv preprint (Tavares et al., 2018) has challenged the conclusions of Rivas et al. (2017). Tavares’ main result comes from using the `--RAFSp` option in our R-scape software. This changes the default G-test statistic to a statistic called RAFS (RNAalifold with stacking), with a background correction called APC (average product correction). Tavares’ conclusions depend on their assumption that RAFS is a covariation measure. However, the RAFS statistic does not solely measure covariation. In (Rivas et al., 2017, online methods and Supplemental Figure 1) we showed controls that led us to conclude that the RAFS statistic, especially when APC-corrected with `--RAFSp`, systematically predicts false positive RNA base pairs on unstructured sequences, making this option unsuitable for distinguishing structural RNA sequence alignments from other conserved sequence alignments. We explain those published conclusions here at more length, using Tavares’ data as additional examples.

While the G-test is a pure covariation measure, the RAFS statistic measures covariation *and consistency*. RAFS assigns relatively high scores to pairs of alignment columns that are consistent with base pairing even if there is no covariation at all. For example, Figure 1 (left) shows an alignment of 20 identical sequences. Tavares’ analysis reports the three proposed base pairs as “significantly covarying” because RAFS sees them as consistently conserved. A conserved G column paired to a conserved pyrimidine (C/U) column has no covariation, but is even more strongly rewarded by RAFS, because it rewards GU/GC half flips. Using RAFS with the APC correction further compounds the problem by (in effect) subtracting the average background signal; scores involving more conserved columns get boosted in a context of less conserved, poorly scoring column pairs.

**Figure 1:**
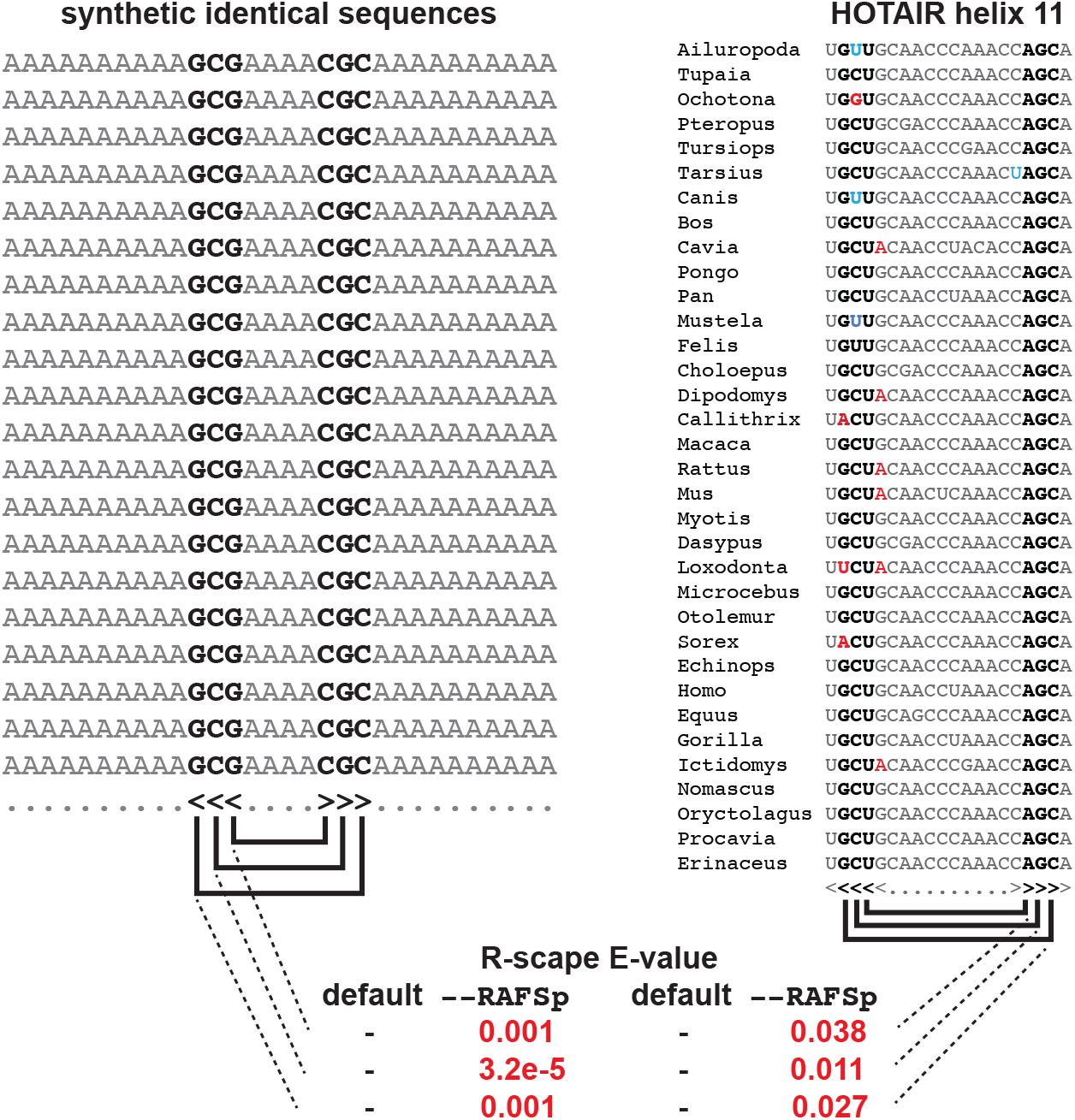
Tavares’ analysis calls pairs of noncovarying conserved primary sequence positions “significantly covarying”. Left: an alignment of 20 identical synthetic sequences with a putative annotated structure of 3 base pairs. Command: `R-scape --rna --RAFSp synthetic.sto`. Right: Tavares’ alignment of HOTAIR domain 1 putative helix 11, showing a five base helix in which they detect “covariation” support for three base pairs (bold). There is no covariation and no compensatory base pair substitutions at these three pairs, by definition, because the AGC on the right side of the proposed three pairs is invariant. Red: substitutions inconsistent with the proposed structure; blue: “half flip” substitutions compatible with GU/GC or AU/GU base pairs.

Figure 1 (right) shows Tavares’ alignment of HOTAIR domain 1 helix 11, where their analysis annotated three significantly covarying pairs in a five base pair proposed helix. The right side of these proposed three base pairs is 100% identical AGC in all sequences, so there is necessarily no covariation and no compensatory base pair substitutions that supports these proposed pairs. Numerous substitutions (red in the Figure) are incompatible with the proposed structure.

A different analysis of the same HOTAIR lncRNA alignments was published by two of Tavares’ authors in Somarowthu et al. (2015). They reach the same conclusion although the two analyses substantially disagree in detail, detecting only partly overlapping sets of “significantly covarying” base pairs. For example, the three proposed base pairs in HOTAIR helix 11 (Figure 1) were reported as conserved or consistent, not covarying, by Somarowthu et al. (2015). In Figure 2, we reproduce Supplemental Figure 4 from Rivas et al. (2017), which we had used to point to related problems in the previous analysis. The 352:370 base pair in proposed HOTAIR domain 1 helix 10 is an illustrative example of where Tavares and Somarowthu et al. (2015) both annotate a significantly covarying pair. The 352 position is a highly conserved G, and the 370 position is a mostly conserved C. There is only one pairwise compensatory substitution (an AU in *Cavia porcellus*), an inconsistent substitution (AC in *Mustela*), and many inconsistent substitutions at other pairs in the proposed helix. The mouse/human pairwise comparison in this region shows six substitutions inconsistent with the proposed structure and no compensatory base pair substitutions.

**Figure 2:**
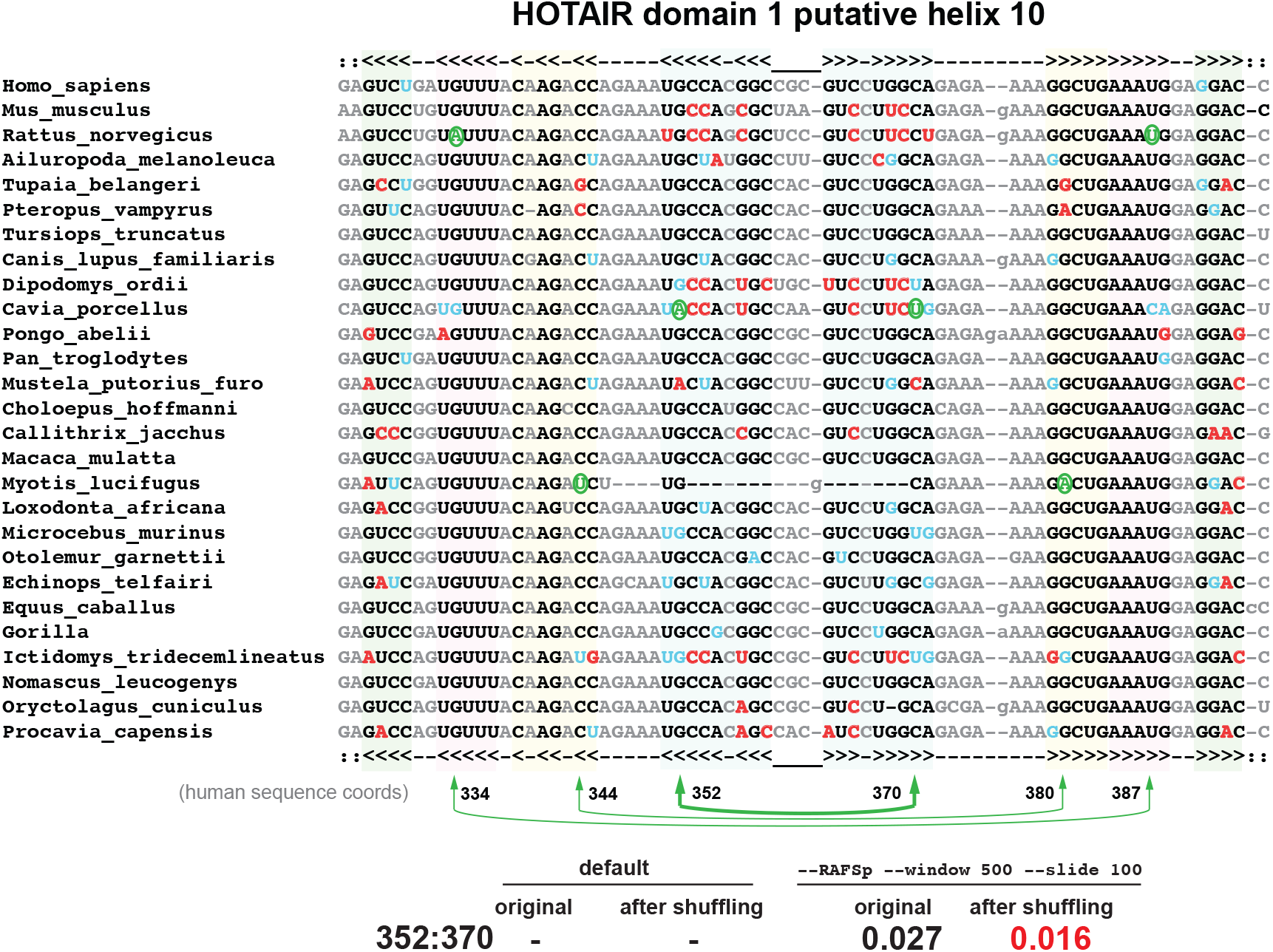
Example of a base pair that both Tavares and Somarowthu et al. (2015) call “significantly covarying” in their HOTAIR D1 lncRNA alignment. The 352:370 pair (in human sequence coords used by Tavares) was called significantly covarying in both Somarowthu et al. (2015) and Tavares; the 334:387 and 344:380 pairs were also called significantly covarying in Somarowthu et al. (2015) but not in Tavares. Green: compensatory base pair substitutions relative to most abundant canonical base pair; blue: “half flips” (such as GC to GU); red: substitutions inconsistent with proposed base pairs. Tavares’ analysis still calls the 352:370 pair significantly covarying even after the residues in each column are permuted to destroy all covariation. Command: `R-scape --RAFSp --window 500 --slide 100 HOTAIR_D1.sto`. Derived from Supplementary Figure 4 of Rivas et al. (2017).

A control for whether Tavares’ analysis is detecting covariation is to shuffle the alignment by permuting the residues in each individual column. This destroys all covariation while preserving position-specific sequence conservation. Figure 2 shows that on a permuted HOTAIR D1 alignment, Tavares’ analysis still calls 352:370 “significantly covarying” (E=0.016). Using Tavares’ `--RAFSp --window 500 --slide` 100 analysis on the complete HOTAIR D1 alignment, similar numbers of “significantly covarying pairs” are detected in permuted alignments (range 28-39, in 10 different shuffles) as in the original alignment (30, at threshold E< 0.05). This is consistent with all their reported signal coming from primary sequence conservation patterns.

The RAFS statistic was developed for the purpose of predicting consensus RNA structures from alignments of sequences already presumed to have a structure (Hofacker et al., 2002; Lindgreen et al., 2006). For this purpose, both covariation and consistency are useful cues. Distinguishing a conserved RNA structure from a conserved primary sequence is a different problem that requires using a statistic that does not systematically detect significant signals on conserved primary sequence alone.

Tavares et al. misstate the conclusions of our paper. We did not conclude that the lncRNAs we analyzed “do not contain conserved structure”. We concluded that based on lncRNA alignments presented in Somarowthu et al. (2015) and other papers, no statistically significant covariation signal of an evolutionarily conserved RNA structure has been detected in these alignments. It is certainly true, as described in our paper (Rivas et al., 2017, Supplemental Figures 1D and 1E, for example) and illustrated by additional examples in Tavares, that statistical power to detect significant pairwise covariation depends on a number of factors, including the accuracy of the alignment (sequence misalignment can destroy true covariations, or even create spurious ones), variability of the individual sites (if positions don’t vary, they can’t covary), the number of sequences in the alignment, and the phylogenetic relationship of the sequences (closely related sequences share spurious correlated variation).

However, a principal conclusion of Somarowthu et al. (2015) was that their HOTAIR lncRNA alignment does suffice to demonstrate “a significant degree of covariation in predicted secondary structural elements” in support of a proposed HOTAIR lncRNA structure. We analyzed exactly the same alignment (Rivas et al., 2017), and so does Tavares. Although there are other data and conclusions in Somarowthu et al. (2015), this particular conclusion remains unsupported and based on a severely flawed analysis, and we believe it should be corrected in the literature.

Tavares et al. describe the covariation analysis in Somarowthu et al. (2015) as “conventional”. In fact that analysis used a data visualization program called R2R that does no statistical analysis, and the R2R authors explicitly warn against interpreting its results for this purpose (Weinberg and Breaker, 2011). R2R only requires a single compensatory pair substitution to annotate a pair as “covarying”, so long as no more than 10%^1^ of the sequences are inconsistent with canonical base pairing of the two positions. (This is why R2R marked the three pairs indicated in Figure 2, for example.) Given some variation and enough sequences, any pair of alignment columns will eventually show a “compensatory pair substitution” by chance. Contrary to Tavares’ statement that R-scape was “developed for application to small, highly structured RNA molecules”, we developed it for any RNA sequence alignment, especially lncRNAs. We were motivated specifically by Somarowthu et al. (2015) and related work, to provide a convenient and statistically sound tool for analyzing RNA sequence alignments for statistically significant base pair covariation signals.

We emphasize that the default analysis in R-scape, a statistic called the G-test, is a conventional pairwise covariation statistic. Similar statistics, such as mutual information, have been used for decades (Gutell et al., 1992, 1994). The principal methodological contribution of R-scape is not the statistic, but rather a computationally efficient means of evaluating statistical significance, using a randomization procedure that preserves the contribution of phylogenetic correlation to false positive covariation signals. R-scape’s randomization procedure does not preserve position-specific primary sequence conservation. This is why RAFSp-detected signals from conserved primary sequence positions can be statistically significant relative to the null hypothesis, in absence of any covariation.

Tavares argue that lncRNAs might pose unusual problems for covariation analysis in part because they are large and might contain only local RNA structures, even though their proposed HOTAIR lncRNA structure is smaller than ribosomal RNA (where covariation analysis was historically instrumental (Gutell et al., 2002)), and the proposed “intricate and modular” HOTAIR secondary structure involves its entire sequence. To make their point, they construct an example where they include a few kilobases of sequence downstream of a conserved SAM-I riboswitch structure element, and claim to show that R-scape can fail to detect a local structure element when it is embedded in a context of long unstructured sequence. This conclusion is incorrect, because besides adding unstructured sequence, Tavares also (perhaps inadvertently) removed the proposed SAM-I secondary structure that they purport to be testing from their alignment data file.

We designed R-scape for two different situations: when a particular secondary structure has been proposed by other means (i.e., you want to test whether any proposed base pairs are supported by significant covariation), or when no structure is yet known (i.e., you want to know if *any* possible base pairs in the alignment are supported, for an as-yet unknown structure). R-scape E-values are corrected for multiple testing, based on the total number of pairs that are considered. Given a proposed structure to test, the “search space” is the number of base pairs in the proposed structure. Without a proposed structure, the search space is all possible pairs. Had Tavares included the proposed SAM-I structure they meant to test, they would have observed that no amount of added unstructured sequence has any effect on R-scape’s power to detect covariations supporting a proposed structure, because the total number of proposed base pairs is unchanged by adding unstructured sequence.

In conclusion, the “improved parameters” reported by Tavares are systematically detecting false positives caused by common primary sequence conservation patterns. Their benchmarks (Tavares, Supplemental Figure 3) showed that the “improved parameters” increase their false positive rate, which they considered to be negligible rather than investigating the source. An E-value is a direct estimate of the expected number of false positives. Any substantial discrepancy between E-values and the observed number of false positives in control experiments should be investigated and accounted for, which Tavares apparently failed to do.

We do not agree that lncRNAs are “well accepted as crucial regulators”, although some specific lncRNAs are. Relatively few lncRNAs have been studied in much detail beyond their expression patterns, and among those that have been studied more closely, there seems to be little coherence in their proposed roles. It seems prudent not to treat “lncRNAs” as a homogeneous class. lncRNAs include a wide variety of different things, including cases where the act of transcription (rather than the RNA) seems to be the effector, cases where an RNA may work by binding proteins or other RNAs, cases of undetected protein-coding mRNAs, and cases where a transcript is nonfunctional “noise” from various sources – none of which needs to involve conserved RNA secondary structure. Sequence analysis should be a powerful means of trying to distinguish any structural lncRNAs from other lncRNAs. Although Tavares refers to our work as “exceedingly stringent”, R-scape’s multiple-test-corrected E-value threshold of 0.05 is only a minimal and standard statistical bar to meet in an exploratory data analysis. We feel that lncRNAs would benefit from more stringent evidence, not less.

## Supporting information

README to expand supplemental material

## Footnotes

1 Somarowthu et al. (2015) customized R2R’s tolerance to allow up to 15% inconsistent base pairs.

