## Supplementary figures and images for "Response to Tavares et al., “Covariation analysis with improved parameters reveals conservation in lncRNA structures”"

### HOTAIR_D1_1.cyk.R2R.sto.pdf

#### HOTAIR_D1_1

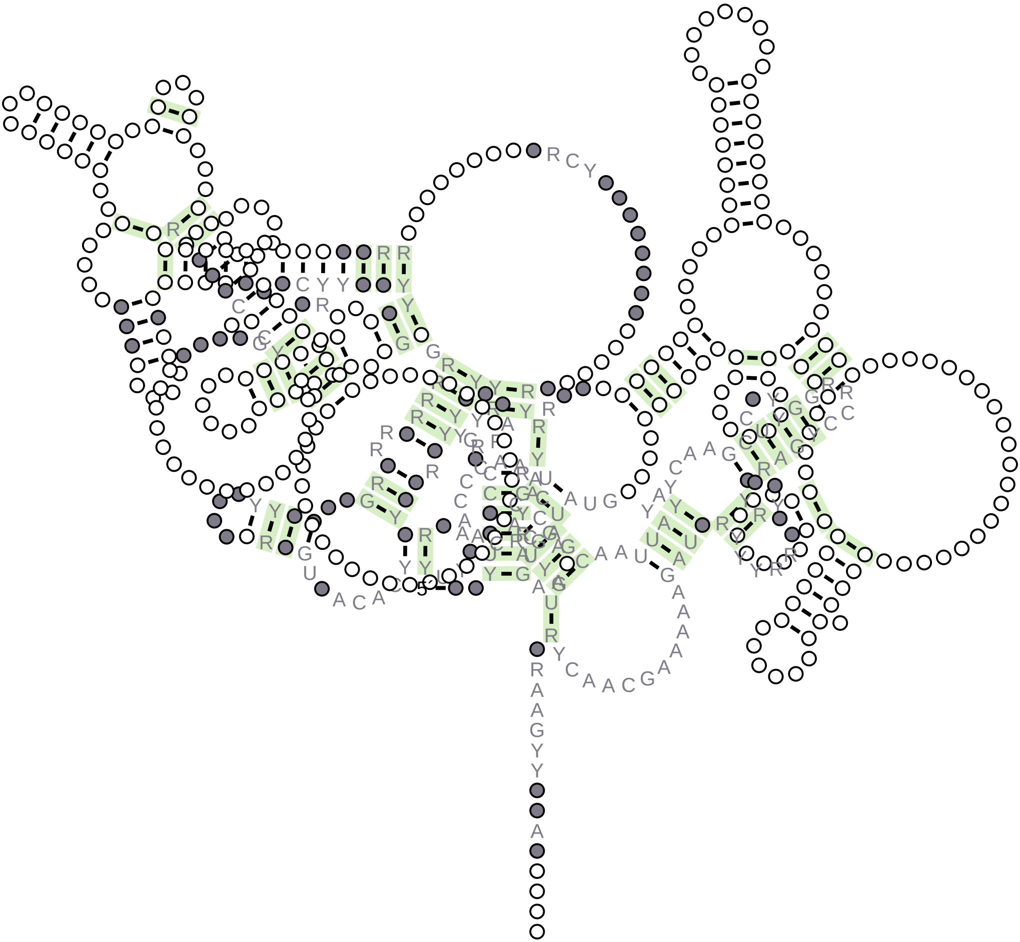

### HOTAIR_D1_1.R2R.sto.pdf

HOTAIR_D1_1

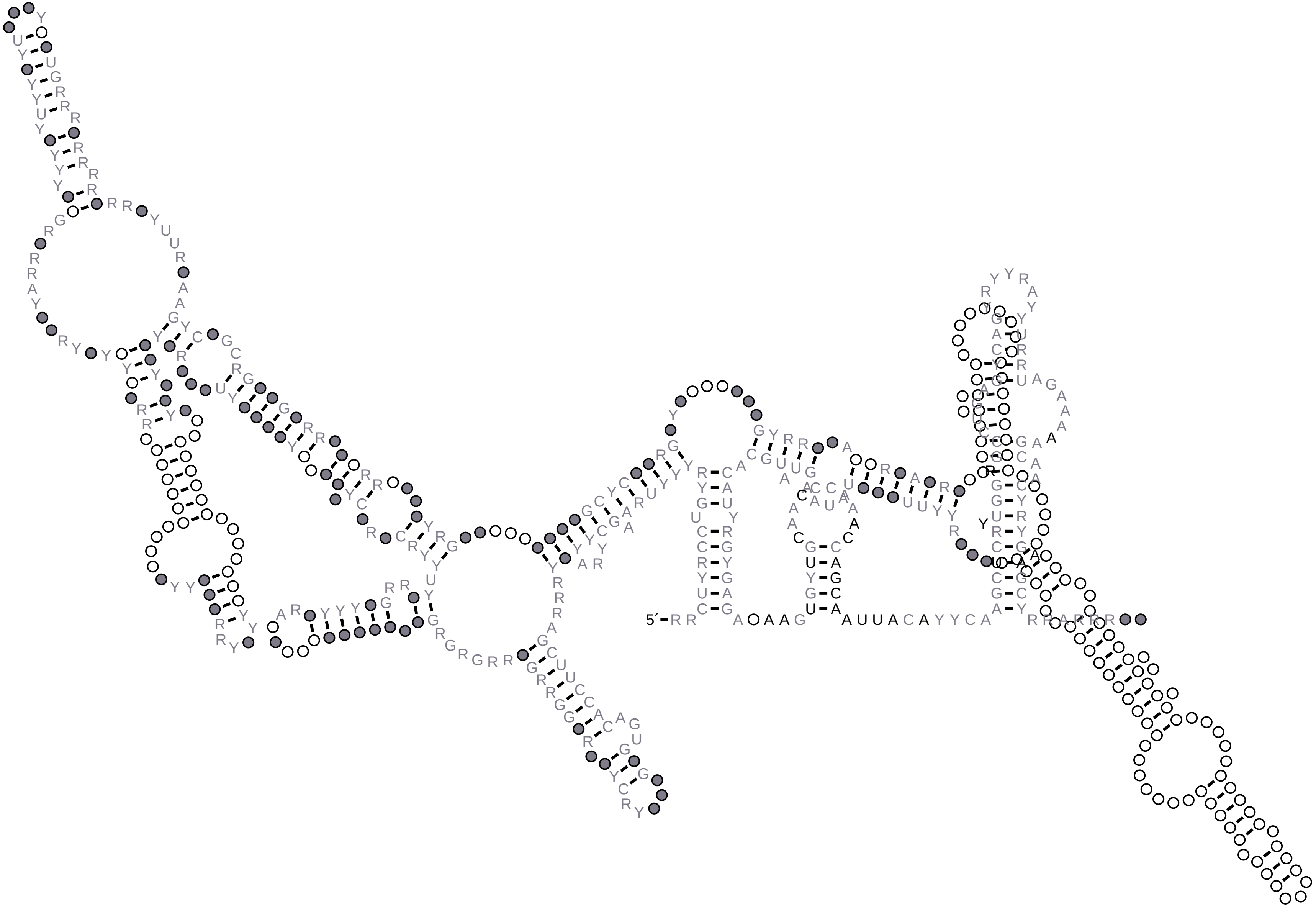

### HOTAIR_D1_1_1-500.R2R.sto.pdf

#### HOTAIR_D1_1_1-500

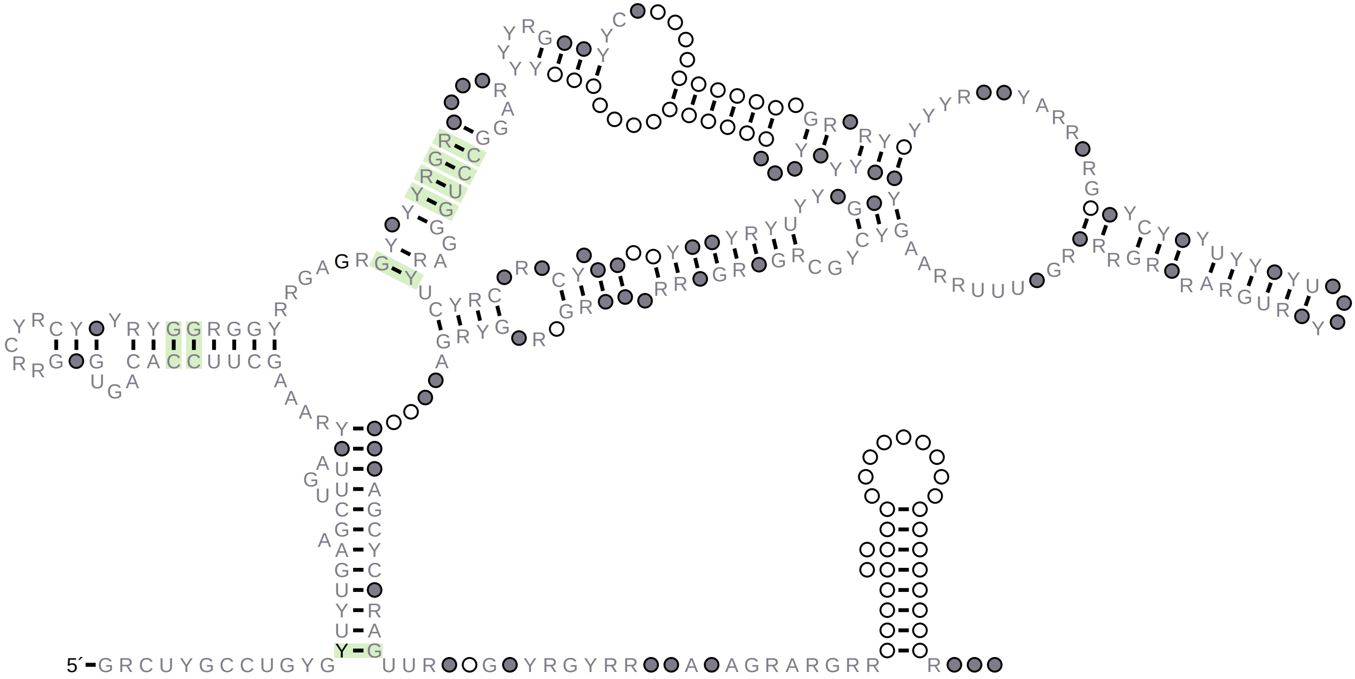

### HOTAIR_D1_1_101-600.R2R.sto.pdf

HOTAIR_D1_1_101-600

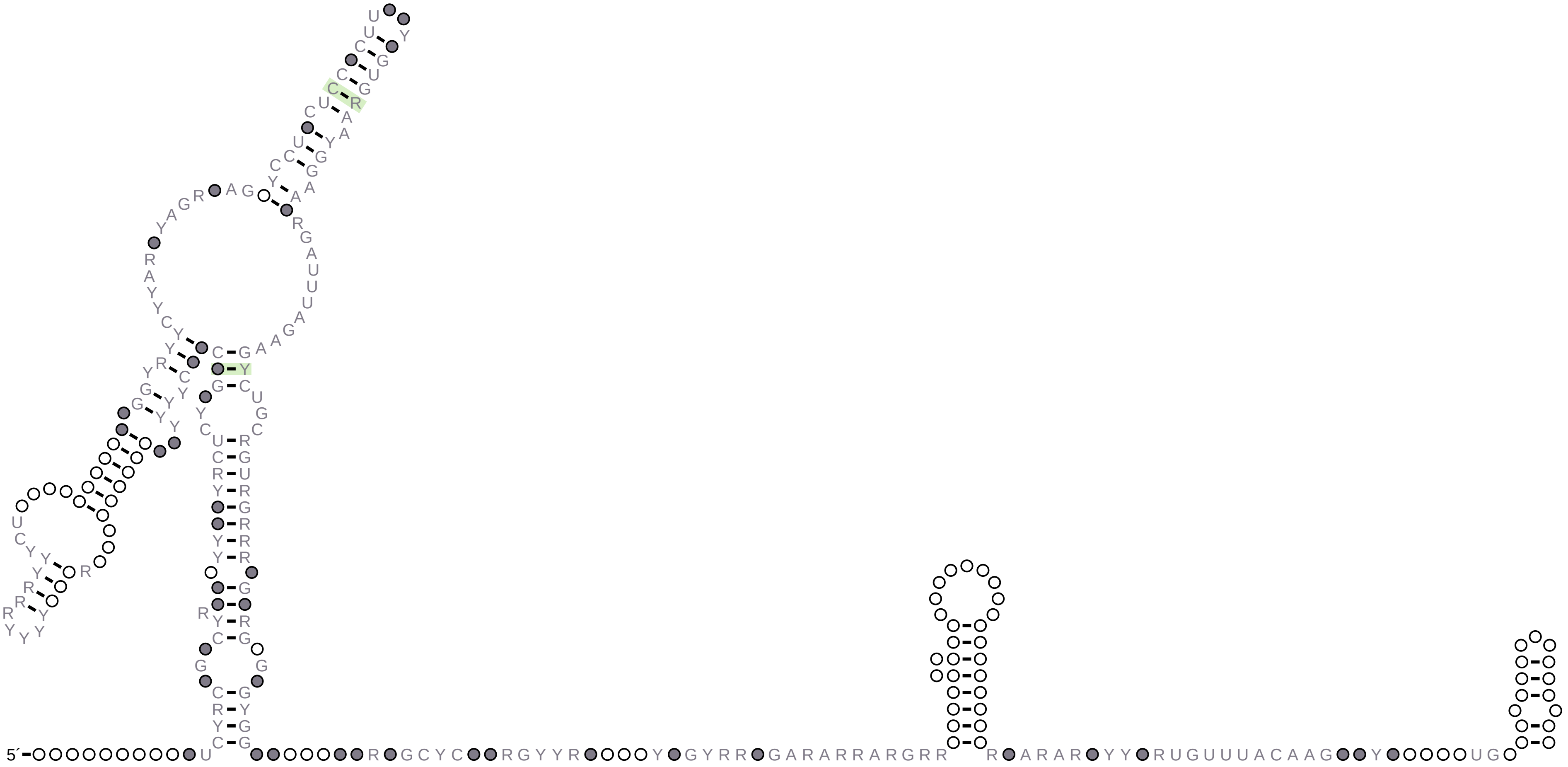

### HOTAIR_D1_1_201-700.R2R.sto.pdf

HOTAIR_D1_1_201-700

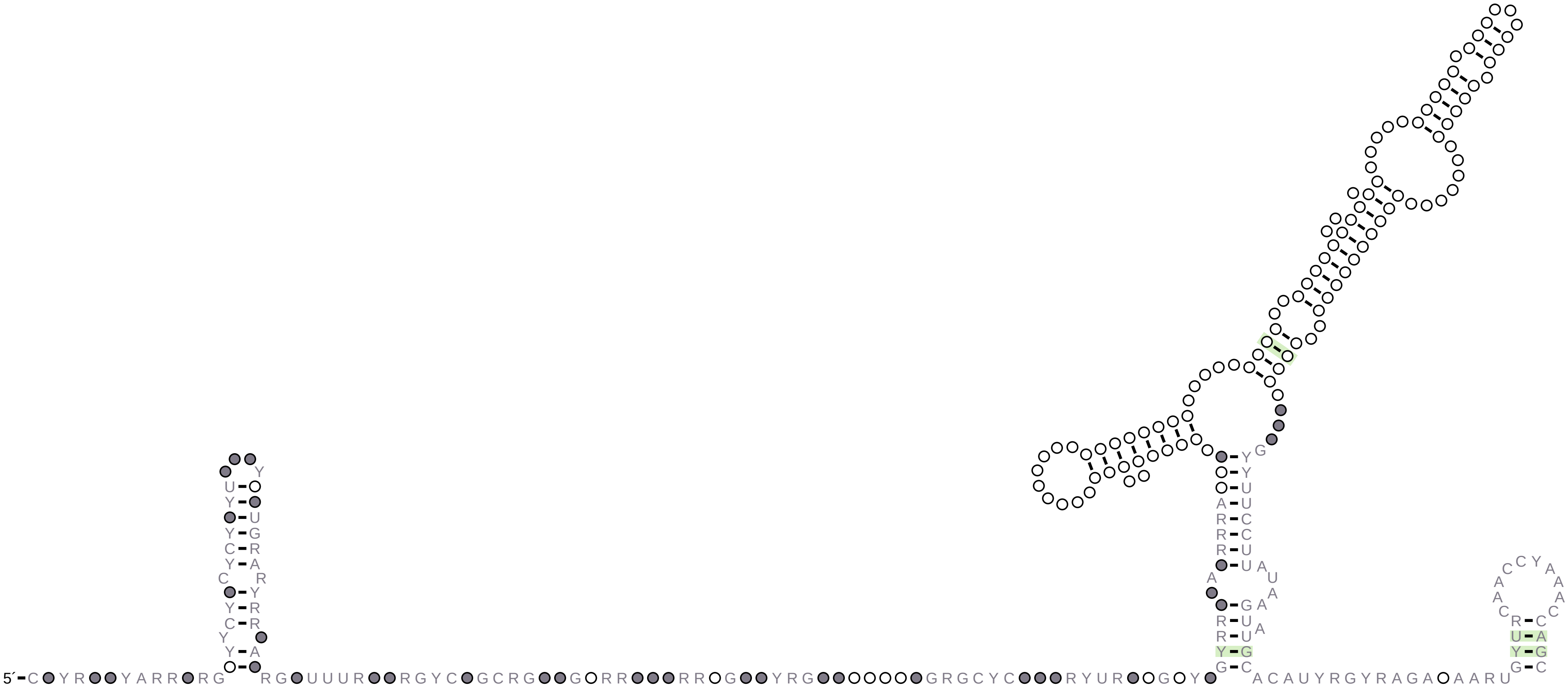

### HOTAIR_D1_1_301-792.R2R.sto.pdf

HOTAIR_D1_1_301-792

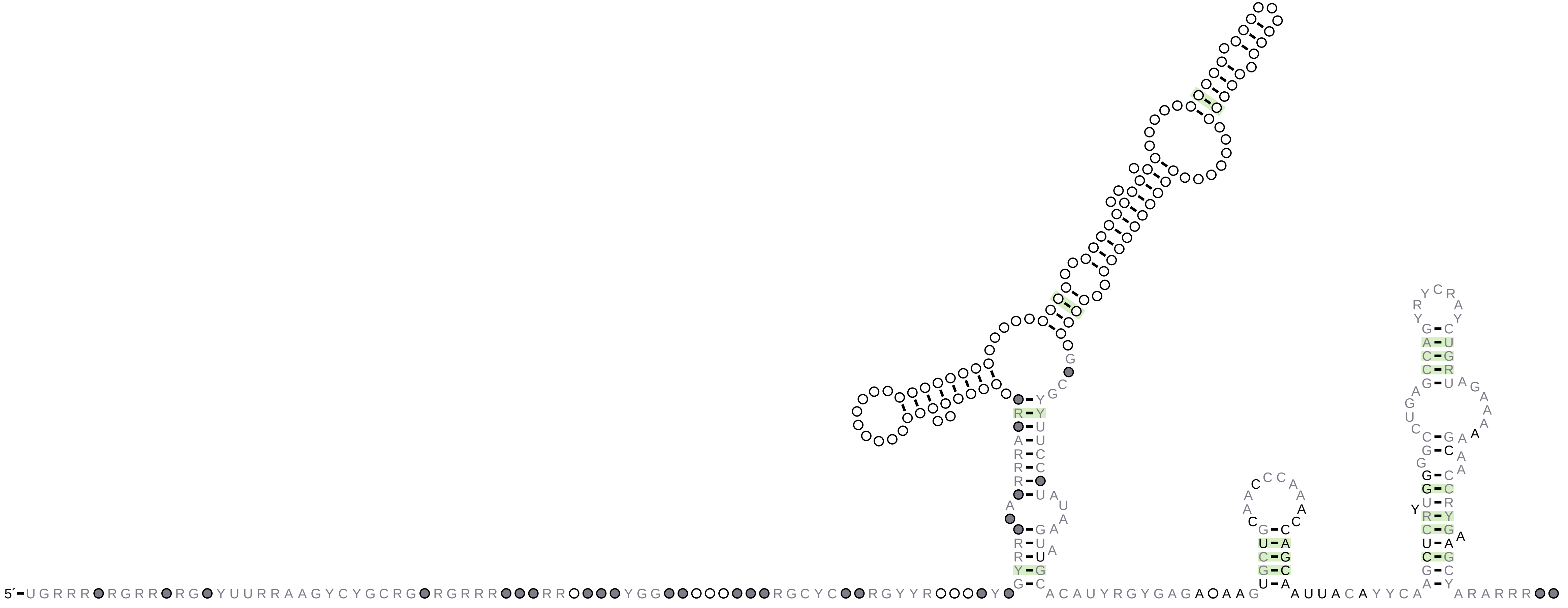

### HOTAIR_H11_1.cyk.R2R.sto.pdf

#### HOTAIR_H11_1

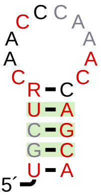

### synthetic_identical_1.R2R.sto.pdf

#### synthetic_identical_1

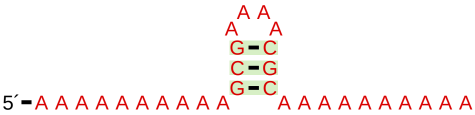
